# Body size and tree species composition determine variation in prey consumption in a forest-inhabiting generalist predator

**DOI:** 10.1101/2020.05.20.105866

**Authors:** Irene M. van Schrojenstein Lantman, Eero J. Vesterinen, Lionel R. Hertzog, An Martel, Kris Verheyen, Luc Lens, Dries Bonte

## Abstract

Trophic interactions may strongly depend on body size and environmental variation, but this prediction has been seldom tested in nature. Many spiders are generalist predators that use webs to intercept flying prey. The size and mesh of orb webs increases with spider size, allowing a more efficient predation on larger prey. We studied to this extent the orb-weaving spider *Araneus diadematus* inhabiting forest fragments differing in edge distance, tree diversity and tree species. These environmental variables are known to correlate to insect composition, richness and abundance. We anticipated these forest characteristics to be a principle driver of prey consumption. We additionally hypothesised them to impact spider size at maturity and expect shifts towards larger prey-size distributions in larger individuals independently from the environmental context.

We quantified spider diet by means of metabarcoding of nearly 1000 *A. diadematus* from a total of 53 forest plots. This approach allowed a massive screening of consumption dynamics in nature, though at the cost of identifying the exact prey identity, as well as their abundance and putative intraspecific variation. Our study confirmed *A. diadematus* as a generalist predator, with more than 300 prey ZOTUs detected in total. At the individual level, we found large spiders to consume fewer different species, but adding larger species to their diet. Tree species composition affected both prey species richness and size in the spider’s diet, although tree diversity per se had no influence on the consumed prey. Edges had an indirect effect on the spider diet as spiders closer to the forest edge were larger and therefore consumed larger prey. We conclude that both intraspecific size variation and tree species composition shape the consumed prey of this generalist predator.

## Introduction

Trophic interactions are a key component of ecological networks (Thébault & Loreau, 2005; Landi *et al*., 2018). Food webs are strongly impacted by the level of diet specialisation of the involved consumers. With higher per capita consumption rates, specialist predators are more effective in prey control than generalists (Diehl, Sereda, Wolters & Birckhofer, 2013), yet opportunistic and generalist predators provide stronger stabilizing effect within a food web (Gross, Rudolg, Levin & Dieckmann, 2009). Generalist species may, however, be a collection of individuals that specialise on specific components of the full prey spectrum (Bolnick *et al*., 2003; Araújo, Bolnick & Layman, 2011). This intraspecific variation is known to arise from behavioural or physiological niche divergence, but equally from environmental variation constraining prey availability and the species spectrum (Bolnick et al. 2013).

Spiders are known to be the principal consumers of insects (Nyffeler & Birkhofer, 2017), and many species are opportunistic and generalist predators that rely on active hunting or trapping for prey capture (Eitzinger *et al*., 2019). Orb-web spiders build vertical webs from silk to intercept flying prey, from small insects to smaller vertebrates (e.g. Brooks, 2012). As interception-feeders, their prey spectrum, is by definition strongly determined by the environment. The diet of generalist and opportunistic predators is therefore expected to reflect the species richness of the prey community (Bison *et al*., 2015; Schmidt, Mosbacher, Eitzinger, Vesterinen & Roslin, T., 2018; Etzinger, Roslin, Vesterinen, Robinson & O’Gorman 2021).

Insect, and thus prey, diversity is largely structured by the plant community composition and plant diversity (Price, 2002; Scherber *et al*., 2010; Rzanny *et al*., 2013). By providing a larger variation in resources, increases in plant diversity are known to promote coexistence of herbivore and predator species (Haddad *et al*., 2009; Hertzog, Meyer, Weisser & Ebeling, 2016; O’Brien *et al*., 2017). Additionally, as plant diversity is linked to an increase in productivity (Loreau *et al*., 2001; Hooper *et al*., 2005), prey abundance and mean prey size should increase as well (Allen *et al*., 2006). Despite the general prediction of plant diversity increasing insect diversity (Scherber *et al*., 2010), plant species identity may have larger effects on the community composition of arthropods than its diversity per se (Vehviläinen, Koricheva & Ruohomäki, 2008; Scherber, Vockenhuber, Stark, Meyer & Tscharntke 2014; van Schrojenstein Lantman et al. 2019). Equally, and independent from these primary producer dynamics, insect and thus spider prey variation can be impacted by the spatial dimensions of their habitat, especially by habitat size and isolation (Debinski & Holt, 2000; Krauss *et al*., 2010). According to the trophic rank hypothesis, higher trophic levels are generally more affected by habitat fragmentation than lower trophic levels (Holt, 2002; Cagnolo *et al*., 2009; Martinson & Fagan, 2014, Hillaert, Vandegehuchte, Hovestadt & Bonte, 2018). Edge densities increase with fragmentation (Haddad *et al*., 2015) and are prominent drivers of insect composition (Murcia, 1995; Schmidt, Jochheim, Kersebaum & Lischeid, 2017) and abundance (Debinski & Holt, 2000; Rand, Tylianakis& Tscharntke, 2006). In forests fragments, arthropod diversity has been shown to be strongly determined by tree species diversity, identify and patch size (Hertzog et al. 2019, 2020; Perring et al. 2021). The patch-size effects are directly translated to edge effects which allow allowing coexistence of both forest and matrix-species (e.g., Rand et al. 2006). The warmer microclimate at forest edges also favours smaller arthropods (Atkinson & Sibly, 1997; Kingsolver & Huey, 2008).

In general, and despite potential variation due to weather conditions and prey availability (Schneider & Vollrath, 1998; Bonte, Lanckacker, Wiersma & Lens, 2008; Sensenig, Agnarsson, Gondek & Blackledge, 2010; Tew, Adamson & Hesselberg, 2015), larger orb web spider species (Dahirel, Dierick, Cock & Bonte, 2017) but also larger individuals of the same species (Bonte *et al*., 2008; Dahirel, De Cock, Vantieghem & Bonte 2019) build larger webs with larger mesh sizes. This allows predation on larger prey that provide essential resources for reproduction (Venner & Casas, 2005). Since predator body size determines the maximum prey size that can be caught (as shown for spiders by Nentwig & Wissel, 1986), large predators can be expected to expand or shift their prey composition towards larger prey (Woodward & Hildrew, 2002). This size-dependency of web-building has been demonstrated in the cross spider *Araneaus diadematus*. As other in species (Dahirel et al. 2017), *A. diadematus* may show individual-level resource specialization in relation to the prey-availability by adapting web building behaviour (Schnieder & Vollrath 1998). More-over, since prey capture does not automatically imply prey consumption (Janetos, 1982), the relationship between spider size and prey (size) consumption remains untested.

Describing and understanding trophic interactions in complex habitats, such as forests, rather than in experiments or simple agricultural systems is a challenging endeavour. We lack a profound understanding on the relative importance of environmental relative to intraspecific variation for realised trophic interactions in nature. We therefore engaged in an unprecedented barcoding (O’Rorke, Lavery & Jeffs, 2012; Pompanon *et al*., 2012) of the species’ gut content to understand to what extent these components of environmental variation as well as predator size determine tropic interactions of *A. diadematus* in forest fragments dominated by different proportions of *Quercus robur* L. (pedunculate oak), *Quercus rubra* L. (red oak) and *F. sylvatica* L. (common beech). We specifically tested the hypothesis that prey richness in the diet increases with both tree diversity, the relative availability of pedunculated oak and forest edge proximity, due to already documented increases in prey species availability (Hertzog et al. 2020). Independent of these tree composition and spatial effects, we tested the prediction that large spiders consume larger prey, and therefore have a wider (size) range of prey species in their diet. Since more diverse forests are expected to contain larger insects, an similar shift in prey consumption is expected under the assumption of orb web spider diet to be primarily determined by prey availability rather than prey selection.

## Materials and methods

### Study site

This study was conducted within the TREEWEB research platform (www.treedivbelgium.ugent.be/pl_treeweb.html) situated in the fragmented landscape of northern Belgium (50.899°N, 3.946°E – 50.998°N, 3.584°E). This platform consists of 53 research plots of 30 × 30 m^2^. All have a similar land-use history (forest since at least 1850), management (no forest management in the last decade) and soil (dry sandy-loam). *Quercus robur* L. (pedunculate oak), *Quercus rubra* L. (red oak) and *Fagus sylvatica* L. (common beech) are the focal tree species in these forests. Plots of the three monocultures and all possible species mixtures (7 different stand compositions in total) were replicated 6 to 8 times along a fragmentation gradient. Edge distance (ranging from 7.0 to 215.5 m) was used as a proxy for edge effects. Edge distance was not correlated to tree diversity, neither did it differ between tree species combinations. Tree diversity was calculated by taking the exponent of the Shannon diversity index. The Shannon diversity index was calculated using the basal stem area of the tree species per plot. For more information on the setup of the study plots see (De Groote *et al*., 2017).

### Study species

We sampled common orb-weaver spiders (*Araneus diadematus* Clerk, 1757) for this study, as they are abundant in the study area and present in all of our study plots. We collected, if possible, 20 adult female *A. diadematus* in each plot from the 29^th^ of August till 8^th^ of September 2016. The spiders were taken from their webs, which were located at breast height. Collected spiders were immediately killed and stored in 99.6% alcohol. In some plots we could not collect 20 spiders, even after multiple visits. Spider size was taken by measuring the maximum width of the cephalothorax (i.e. carapace) under a stereomicroscope using a calibrated eyepiece graticule. Cephalothorax or carapace width has been a common used proxy for body size in mature spiders (Hagstrum, 1971; Greenstone, Morgan & Hultsch, 1985).

### Molecular analysis

To establish the diet of the spiders, we used a proven metabarcoding protocol for spiders and other invertebrate predators (Vesterinen, Lilley, Laine & Wahlberg 2013; Kaunisto, Roslin, Sääksjärvi & Vesterinen, 2017; Eitzinger *et al*., 2019). Shortly, we extracted DNA from the spiders’ abdomen using NucleoSpin ® Tissue kit (cat. nr. 740952.250, Germany). To amplify mitochondrial COI gene, we used primers ZBJ-ArtF1c and ZBJ-ArtR2c from (Zeale, Butlin, Barker, Lees & Jones, 2011). As these primers also amplify the spiders themselves, we designed a blocking primer to decrease predator amplification in favour of prey amplification (Vestheim & Jarman, 2008). To prepare the blocking primer, we first downloaded all unique *A. diadematus* sequences from BOLD and GenBank and aligned them with multiple potential prey sequences using Geneious (Kearse *et al*., 2012). Then we designed three primer sequences that overlapped the reverse primer ZBJ-ArtR2c and that were specific for *A. diadematus* (zero mismatches) but that did not match to any potential prey. The blocking primers were tested using primer BLAST (Koressaar & Remm, 2007; Untergasser *et al*., 2012; Ye, Coulouris, Zaretskaya, Cutcutache, Rozen & Madden, 2012). The best candidate (that did not bind to anything in the database except *A. diadematus*) was chosen. This primer sequence was ordered with C3 spacer modification at the 3’ end (Aradia-R-blk-C3: 5’-CCA AAT CCC CCA ATT AAA ATA GGT ATA-C3 spacer -3’). PCR conditions and library preparation followed (Kaunisto *et al*., 2017) and (Vesterinen, Lilley, Puisto & Blomberg, 2018). Shortly, the first PCR included locus-specific linker-tagged primers, and in this stage the target gene was amplified. In the second PCR, we introduced linker-tagged indexed (different index in forward and reverse primers) adapters that were compatible with Illumina platforms and perfectly matched the linker-tags in the first initial PCR. To minimize the risk of contamination, all the extraction steps were carried out in carefully cleaned lab space, using purified pipettes with filter tips. All the extraction batches included negative controls to account for contamination issues. Washing the spiders several times in 99.6% ethanol during the collection, storage and preparation for extraction process was deemed to be appropriate for sterilization. Negative controls containing all but template DNA were included in each PCR assay. PCR products were never introduced to the pre-PCR space. All the uniquely dual-indexed reactions were pooled and purified using SPRI beads as in (Vesterinen *et al*., 2016). The pool in this study was combined with a similarly prepared pool (from a vertebrate dietary analysis) to increase nucleotide diversity in the sequencing and to lower costs per project. Sequencing was performed by Macrogen Korea (Macrogen Inc., Seoul, Rep. of Korea) using HiSeq4000 with TruSeq 3000 4000 SBS Kit v3 chemistry and 151 bp paired-end read length following HiSeq 3000 4000 System User Guide (Document #15066496 v04 HCS 3.3.52).

After sequencing, the reads separated by each original sample were uploaded on CSC servers (IT Center for Science, www.csc.fi) for trimming and further analysis. Trimming and quality control of the sequences were carried out as in (Vesterinen *et al*., 2018). Briefly, paired-end reads were merged, trimmed, and collapsed using 64-bit software VSEARCH (Rognes, Flouri, Nichols, Quince & Mahé, 2016). For chimera-filtering, denoising, and clustering into ZOTUs (‘zero-radius OTU’), we used 32-bit USEARCH (Edgar, 2010; Edgar & Flyvbjerg, 2015). Before collapsing, primers were removed using software Cutadapt (M_ARTIN_, 2011). Then, ZOTUs were mapped back to the original trimmed reads using VSEARCH, and finally ZOTUs were assigned to prey taxa as explained below.

### Data preparation

We summed the presence or absence of each prey taxon in each sample to end up with a frequency of occurrence (FOO) for each prey taxon. Additionally, all the frequencies were scaled to per cent of occurrence as explained in (Deagle *et al*., 2019), creating a modified frequency of occurrence (MFO). We identified prey to the species level, where possible. The ZOTUs were initially identified using local BLAST against all COI sequences downloaded from BOLD and GenBank (Altschul *et al*., 1990; Ratnasingham & Hebert, 2007). When species name was not available but match to the database was high, we used BIN codes from BOLD (Ratnasingham & Hebert, 2013). For details of the ZOTUs see Supplementary materials 1. To visualize the trophic interactions structures resolved by the molecular data, we used package BIPARTITE (Dormann, Fruend, Bluetghen & Gruber, 2008; Dormann, Grubver & Fruend 2009) implemented in program R (R Core Team, 2018). Semi-quantitative foodwebs were constructed using per cent of occurrence as explained above.

For further analysis, the cut-off threshold per ZOTU for the number of reads was set 0.05% of the total number of reads per spider, with a minimum cut-off threshold of 10 reads. A first multivariate analysis was performed to explore the variation in prey composition within the spider diet. The variation in the prey ZOTU composition within the diet of individual spiders was analysed as a function of tree species combination, edge distance and spider size. A distance-based redundancy analysis (Euclidean distance) using the CAPSCALE function from the RDA package (Guo *et al*., 2018) was applied. We performed an analysis of variance on the distance-based redundancy analysis with 1000 permutations (permanova) to quantify the variation in prey species composition explained by the different variables. Taxonomic units (prey species) were treated as binary data (absence or presence) as the data collection and chosen set-up using the metabarcoding technique does not allow a more quantitative approach. We detected DNA from potential parasitoids (e.g. Tachinid flies, Ichneumonid wasps) as well. These records were rare, and can be equally part of the diet. The metabarcoding does, in the same vein, not allow separating potential DNA from species consumed by predators that are here predated by spiders (e.g., Vespidae, Staphylidae). If some bias is created by these, we don’t expect this to be associated with specific environments or size classes.

### Response variables

Per individual spider with known size, four diet-related response variables were calculated. Prey richness was taken to be the number of ZOTUs in the diet of each spider. For every single assigned prey ZOTU, its size (body length) was taken from literature (Supplementary materials 1). Prey size was taken to be the average prey size of taxonomic units preyed by each spider. These two response variables allow us to answer the four specific hypothesis. In addition, analyses were performed with response variables to unearth possible explanations for the found patterns. Taxonomic units with a body length over 1 cm are considered to be of highest gain (Venner & Casas, 2005), and could help us understand patterns arising. Prey richness of large prey was the number of taxonomic units with a body length of over 1 cm present in the diet of each spider. Prey size of large prey was the average prey size of taxonomic units larger than 1 cm present in the diet of each spider. Additional to the four individual-level response variables, we additionally calculated the Sørensen index (within-plot turnover of species composition in the diet), coefficient of variation (CV) for prey richness and CV for prey size were calculated at plot level.

### Statistics and model selection

Four models were applied to the response variables to explore different aspects of the data. M_div_ was the first model in which we tested for the effects of spider size, edge distance, tree diversity and the interacting effects of edge distance and tree diversity. The three other models were a set of models to compare to each other in order to understand the effects of spider size, edge distance and tree species composition (Kirwan *et al*., 2009). The null model (M_null_) includes only spider size and edge distance and assumes no effect of tree species composition. The additive model (M_add_) includes, besides spider size and edge distance the relative basal area of each of the three focal tree species, and the intercept was forced through zero. This model assumes that tree species exert only additive identity effects. The pair-wise interaction model (M_pair_) includes additionally the pair-wise interactions between the relative basal areas of the focal tree species. This model assumes not only additive effects, but also interacting effects between the tree species. The three composition models (M_null_, M_add_ and M_pair_) were compared to each other to understand in which way the tree species composition impacted on the diet-related response variables. The model with the lowest AICc (obtained using the AICcmodavg package by (Mazerolle, 2017) was considered the best. Spider size, edge distance and tree diversity were scaled around their mean in all models (Schielzeth, 2010). Spider size was excluded as an explanatory variable from the models applied to spider size itself. For the plot-level mean response variables, an average plot-level spider size was included as a variable, instead of individual-level spider size.

All analyses were performed in R, version 3.5.1 (R Core Team, 2018). All models, except the models with Sørensen index and coefficients of variation as a response variables, included plot ID as a random factor. This accounts for our data structure and the potential effect of plot ID. Models with overall prey size as response variable had a negative binomial distribution (log-link) with a variance increasing quadratically to the mean, applied through the GLMMTMB package (Brooks *et al*., 2017). Models with the overall prey richness as response variable had a negative binomial distribution with constant variance using glm. Models with the richness of prey larger than 1 cm as response variable had a Poisson distribution. All other models had Gaussian distributions. All models are two-tailed and have no effect of the explanatory variables as H_0_.

## Results

### General results

A total of 340 distinct prey species from 85 families in 8 orders and two classes were detected (Figure 1). The HiSeq 4000 sequencing yielded 265 871 470 paired-end reads. After assigning these reads to unique dual-indexes used in this study, and after trimming and filtering, we ended up with 5765096 prey reads that could be mapped to the original samples. A total of 983 spiders were included in the molecular analysis, and 857 (87.2%) of these provided prey data after bioinformatic filtering including 298 prey species and were included in the subsequent analysis. The highest observed prey species richness within a single spider sample was 15 prey ZOTUs. The most frequent prey detected was *Phaonia pallida* (N = 357), a forest-living muscid fly. The average prey size was 7.5 mm (SD ± 4.1), with only 39 prey species or genera larger than 1 cm. For a full list of prey taxa, and size data, see Supplementary materials 1.

**Figure 1.**
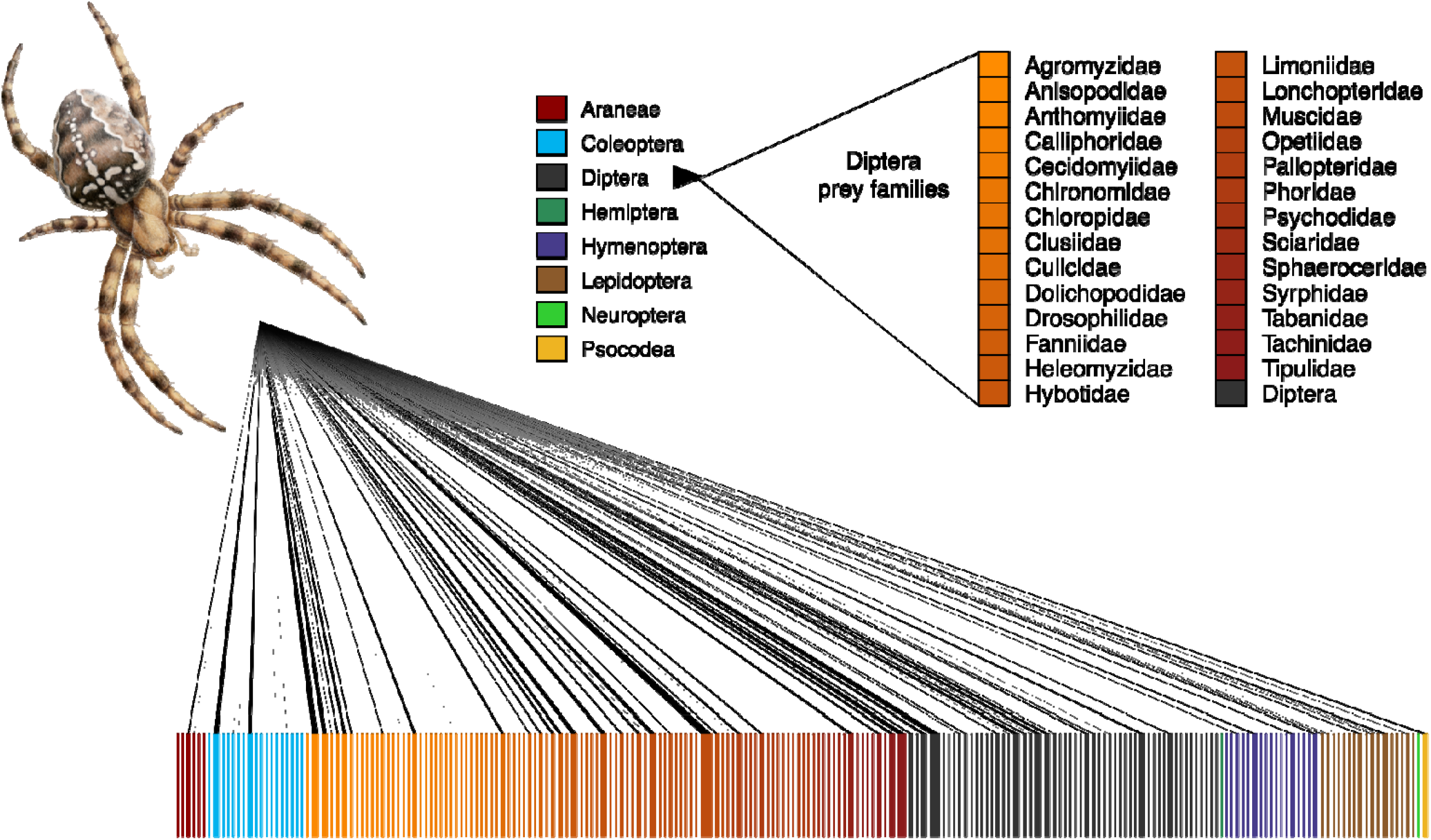
Visual representation of the taxonomic distribution and quantified strength of trophic links from *Araneus diadematus* to their prey. The blocks in the lower row represent prey species. A line connecting the predator with a prey represents detected predation events, and the thickness of the line represents the modified frequency of occurrences (MFO) of each predation record. See the “Data analysis” in the main text for details on the MFO.

Spider cephalothorax width ranged from 2.6 to 5.31 mm (average of 3.57 mm). Spider size decreased further from the edge; tree species identity had additive effects on spider size (Table 1-3, Figure 2). Spider size was not impacted by tree diversity or the interaction between tree diversity and edge distance, but size was largest in monocultures of *Q. robur* and smallest in monocultures of *F. sylvatica* (Table 1-3, Figure 2).

**Table 1.**
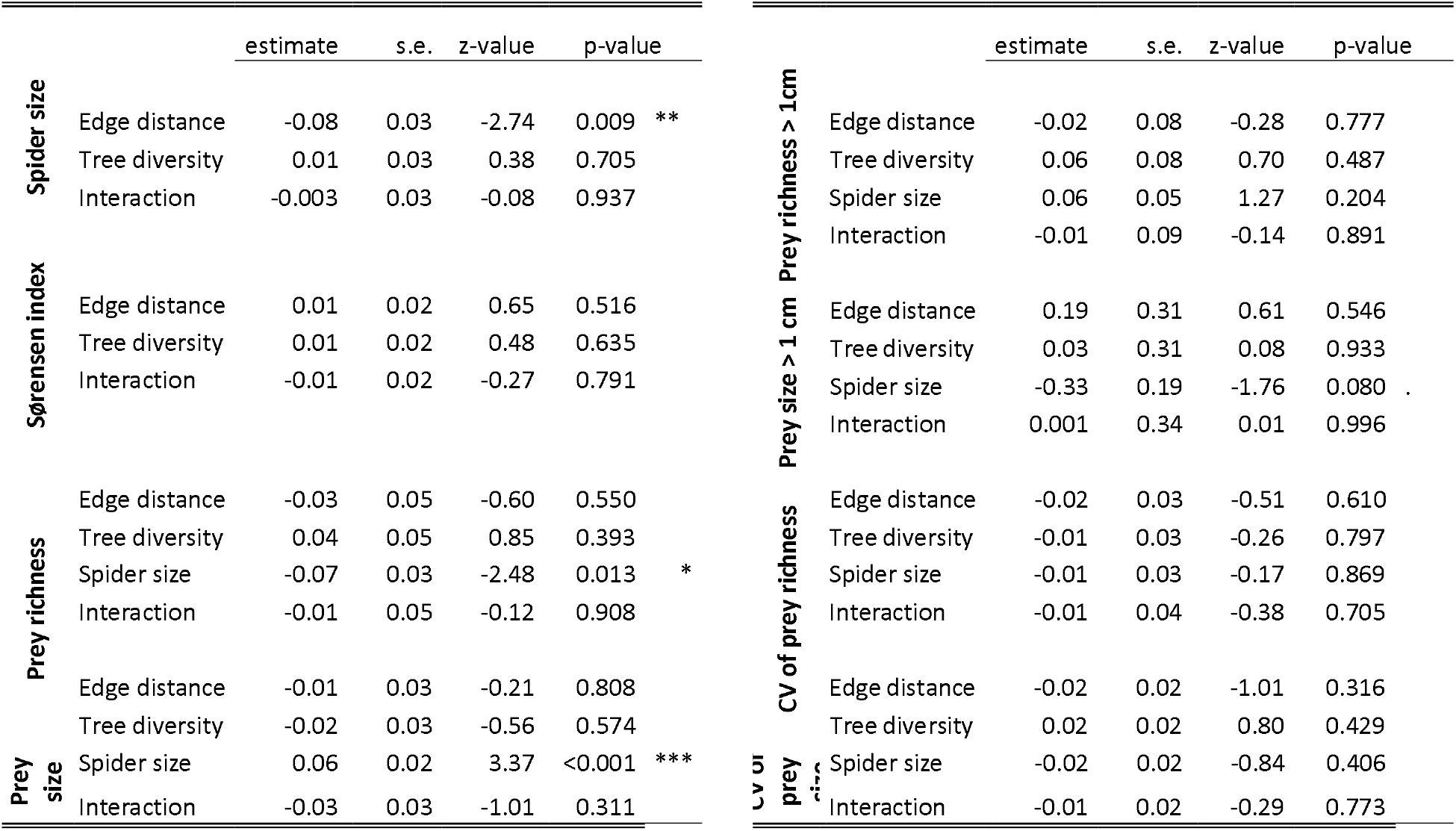
Summary of results for all M_div_ models. Except for the Sørensen index, Plot ID was added as a random factor in the models, but is not shown here. CV stands for coefficient of variation. (. p < 0.100; * p < 0.050, ** p < 0.010, *** p < 0.001)

**Table 2.** Comparison of the tree species composition models for all response variables. Best fitting models based on AICc and parsimony are bold. CV stands for coefficient of variation.

|  | AICc |  |  |
| --- | --- | --- | --- |
|  | M<sub>null</sub> | M<sub>add</sub> | M<sub>pair</sub> |
| Spider size | 1169.26 | **1164.76** | **1164.70** |
| Sørensen index | **-72.46** | -68.54 | -62.71 |
| Prey richness | **4169.50** | 4173.21 | 4175.28 |
| Prey size | 4521.33 | **4512.74** | 4517.46 |
| CV of prey richness | **-0.94** | 3.38 | 6.37 |
| CV op prey size | -58.96 | **-61.93** | -56.76 |
| Prey richness > 1cm | **1866.22** | **1864.35** | **1865.08** |
| Prey size > 1cm | 2173.30 | 2173.76 | **2161.18** |

**Table 3.**
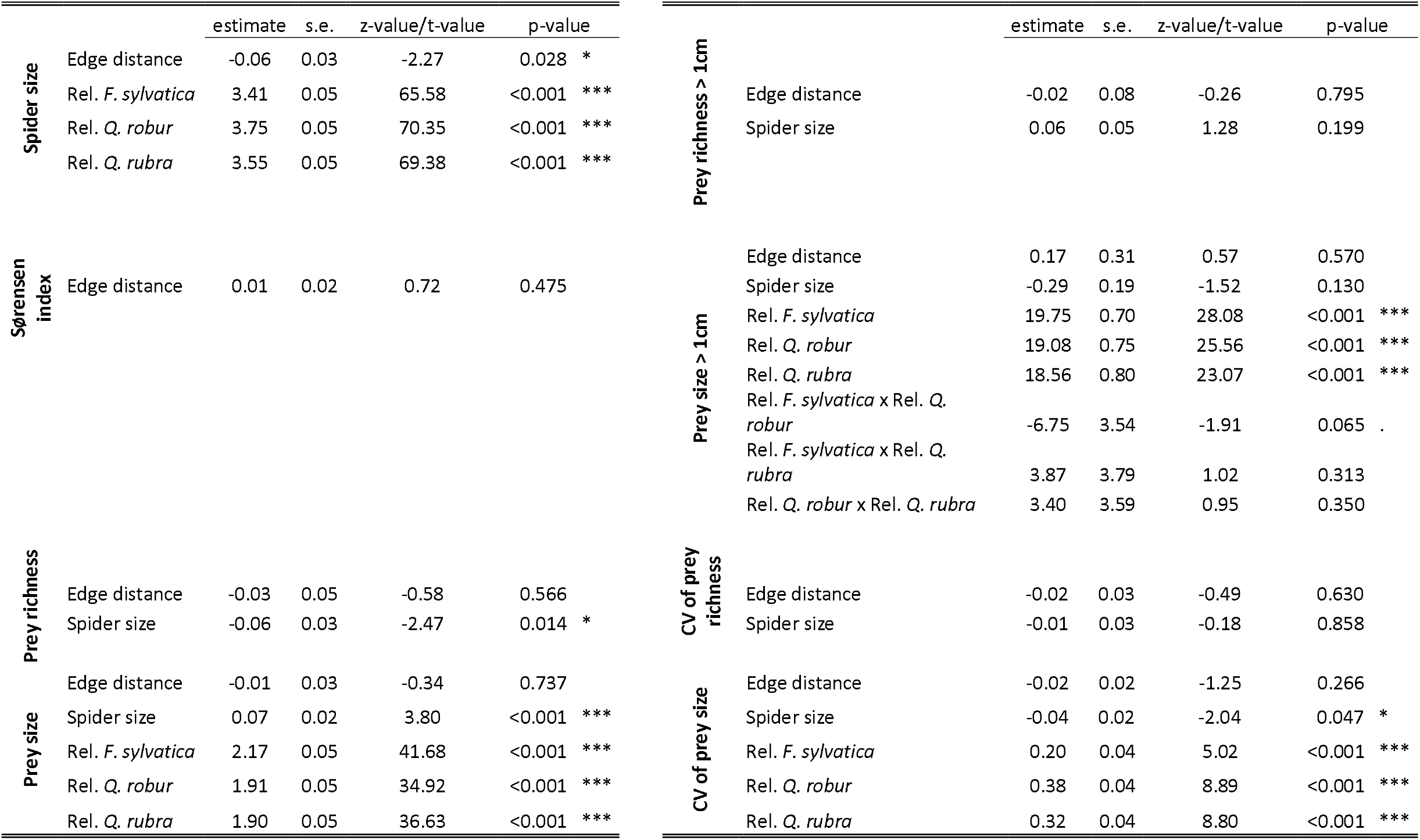
Summary of results for the best fitting models which test for tree species composition effects. CV stands for coefficient of variation. (. p < 0.100; * p < 0.050, ** p < 0.010, *** p < 0.001)

**Figure 2.**
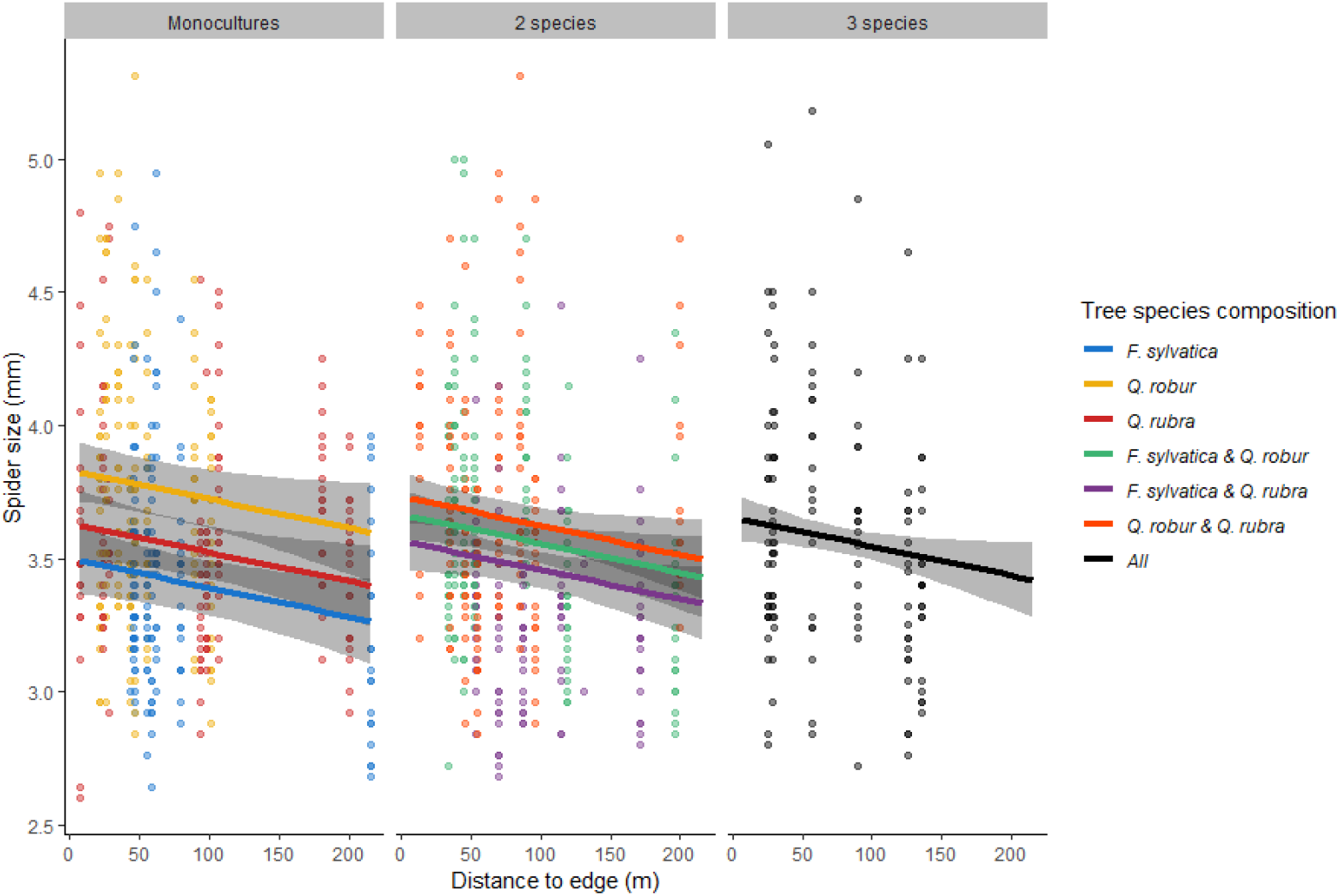
Relationship between the spider’s cephalothorax size (mm) and distance to the forest edge (m) per tree species composition. Data points are the individual spiders (N = 983). Lines with 95% CI are the estimated slope based on model M_add_. Colours refer to the tree species composition. There is a negative relationship between spider size and edge distance, and additive tree species composition effects, where mixtures are averages of each individual monoculture contributing to the mixture.

### Diet composition

The composition of prey species in spider diets is highly variable (Figure 3). Although the constrained components of the ordination only explained 2,2% of the variation in composition, spider size (PERMANOVA, F_pseudo_ = 4.41, p = 0.001) and tree species combination (PERMANOVA, F_pseudo_ = 2.65, p = 0.001) strongly influenced prey species composition. However, all tree species compositions overlapped and showed large variation (Figure 3). Edge distance had no effect (PERMANOVA, F_pseudo_ = 1.28, p = 0.127). Compositional similarity of spider diet as measured by the Sørensen Index was not correlated to edge distance, tree diversity or tree species composition (Table 1, 3).

**Figure 3.**
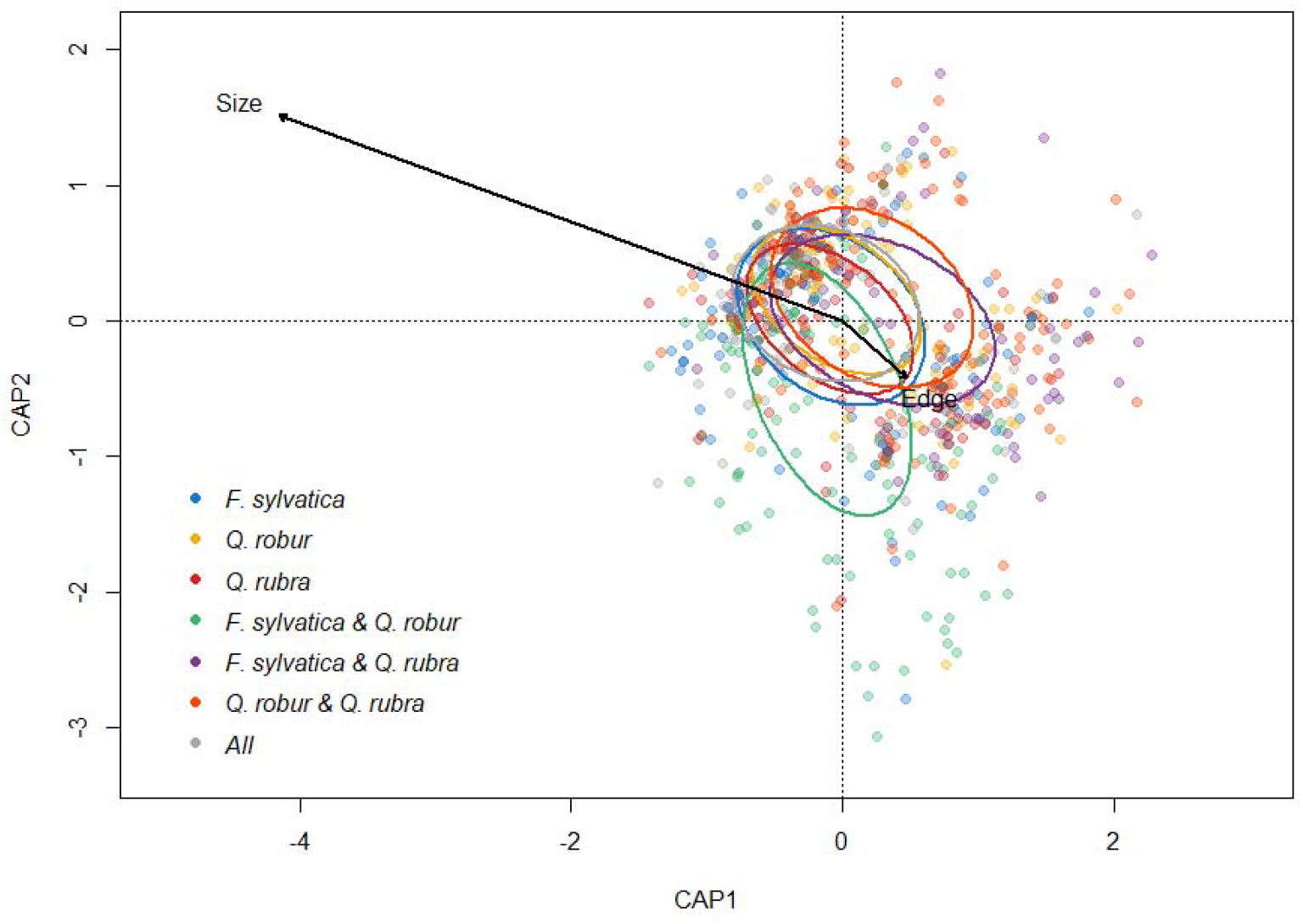
Distance-based redundancy analysis on the prey composition of individual spiders. Colours refer to the different tree species compositions in which the spiders were sampled. Ellipses correspond with the standard deviation of each tree species composition centroid. Size refers to the spider size and Edge to edge distance. The constrained components of the ordination (spider size, edge distance and tree species composition) explained 2.2% of all variation in diet composition.

### Prey richness & size

The models including tree diversity (M_div_) revealed that edge distance, tree diversity and the interaction between them had no impact on the four diet-related response variables: prey richness, prey size, richness of prey > 1cm and size of prey > 1cm (Table 1). In the best fitting composition models (M_null_, M_add_, M_pair_) edge distance had no impact on the diet-related response variables either (Table 3). Spider size had an impact on the overall prey richness and prey size, only marginally on the size of prey larger than 1 cm, and no impact on the richness of prey larger than 1 cm (Table 1). A lower prey richness was detected in large spiders (Figure 4), although prey were larger in these spiders. The effect of identity of the tree species and their relative contribution to prey richness was absent, as M_null_ was the best fitting model (Table 2). For prey size there were additive effects of tree identity (Table 2). Spiders in monocultures of *F. sylvatica* consumed the largest prey species, whilst spiders from *Q. robur* and *Q. rubra* monocultures had very similar sized prey (Table 3, Figure 5). Spiders inhabiting mixtures consumed prey which were the average size of the prey in the monoculture values of tree species included in the tree species composition.

**Figure 4.**
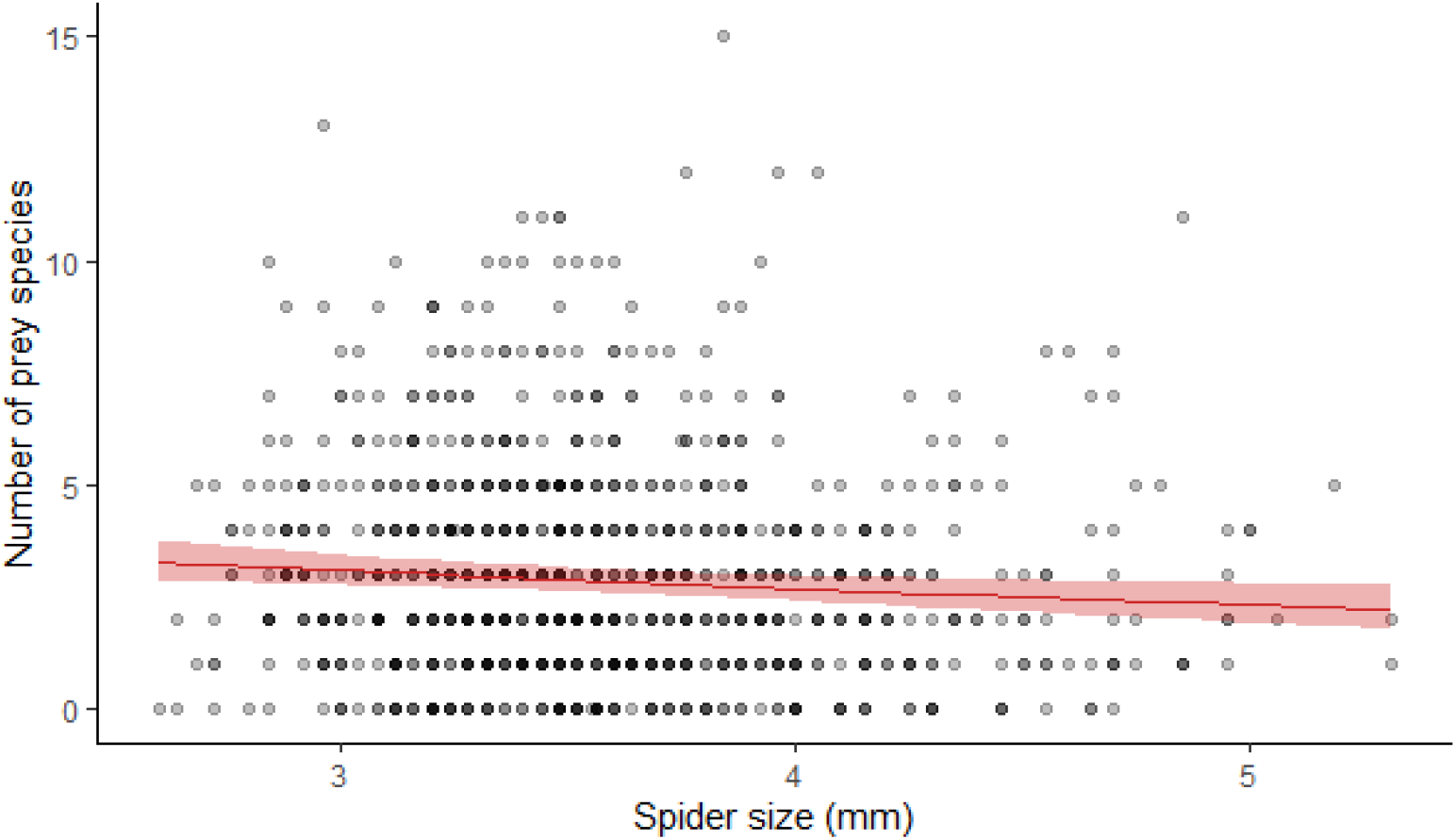
Relationship between the number of prey species in the diet of a spider and its cephalothorax size in mm. Data points are the number of prey species per individual spider (n = 983). Red line with 95% CI is the estimated slope based on model M_div_

**Figure 5.**
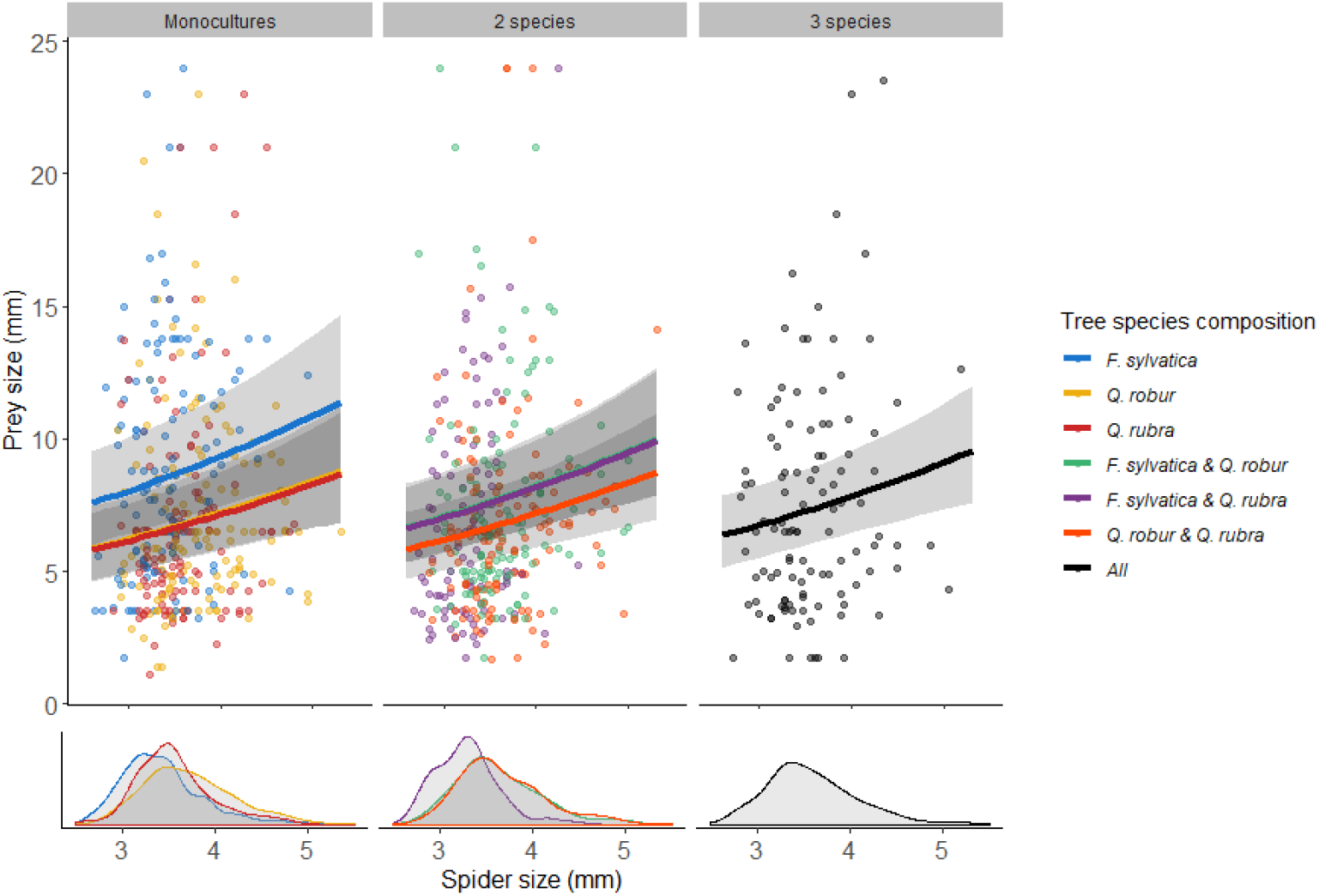
Relationship between the average prey size (mm) of the species caught and the spider’s cephalothorax size (mm) per tree species composition. Data points are the individual spiders (N = 983). Lines with 95% CI are the estimated slopes based on model M_add_. For estimation, edge distance was taken to be the overall average. Colours refer to the tree species composition. There is a positive relationship between average prey size and spider, and additive tree species composition effects, in which monocultures of *F. sylvatica* catch relatively larger prey species than the other monocultures. In mixtures, the prey size are averages of each individual monoculture contributing to the mixture. The lower part of the graph shows density plots of the spider size distribution within each tree species composition.

Richness of prey with the highest gain (prey larger than 1 cm) showed no effects of tree species identity; size of prey with the highest gain (prey larger than 1 cm) did show effects of tree species identity (Table 1-3). These large prey were proportionally most abundant in the diet of spiders from *F. sylvatica* monocultures (Figure 6A). Prey body size was lowest in *F. sylvatica* - *Q. robur* mixtures, relative to their anticipated size in the respective monocultures (Table 3, Figure 6B).

**Figure 6.**
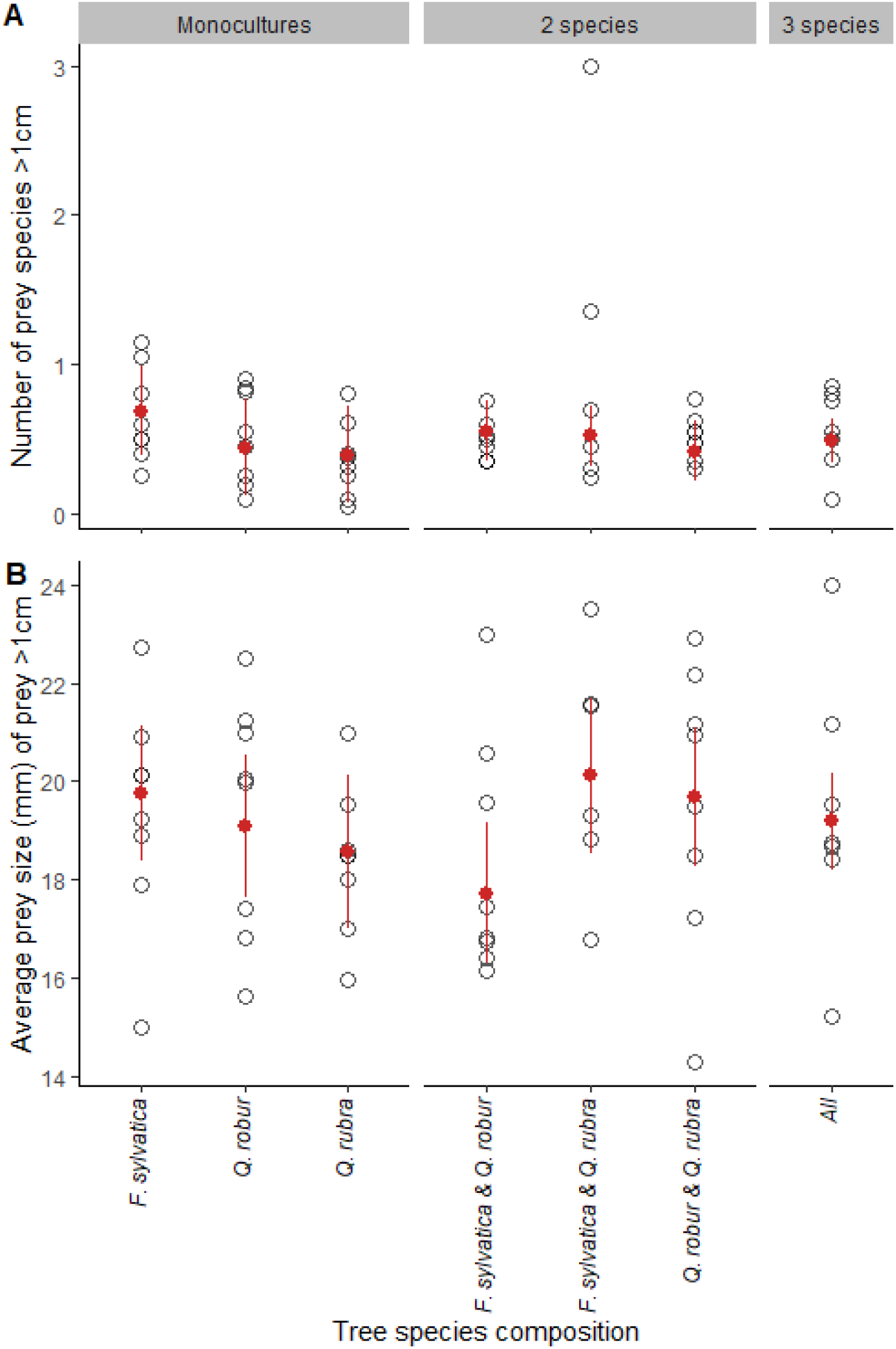
Tree species composition effect on (A) the number of prey species larger than 1 cm and (B) the average prey size of prey larger than 1 cm. Open data points are the averages per plot. Red point ranges are the estimated values based on the best parsimonious model. Edge distance and spider size were taken to be the overall average. For (A) there are additive effects of the monocultures in mixtures, averaging out the number of prey species. For (B) there are pairwise interaction effects in combining *F. sylvatica* with *Q. robur* results in (marginally) lower prey size than expected based on the monocultures.

### Patch-level individual variation in prey consumption

The coefficient of variation (CV) for prey richness revealed that across tree diversity, tree species composition, edge distance and spider size, the level of specialization was the same, as none of the explanatory variables were significant (Table 1), and M_null_ was the best fitting composition model (Table 2). The CV for prey size revealed that plots with on average larger spiders were more consistent in their consumed prey size, independent of tree diversity or edge distance (Table 1). Spiders in monocultures of *Q. robur* were least consistent in the consumed prey size, while those in *F. sylvatica* monoculture were most consistent (Table 3, Figure 7). Spiders inhabiting mixtures showed a level of consistency in prey size that was the average of the monoculture values of tree species included in the tree species composition (Table 1; Figure 7).

**Figure 7.**
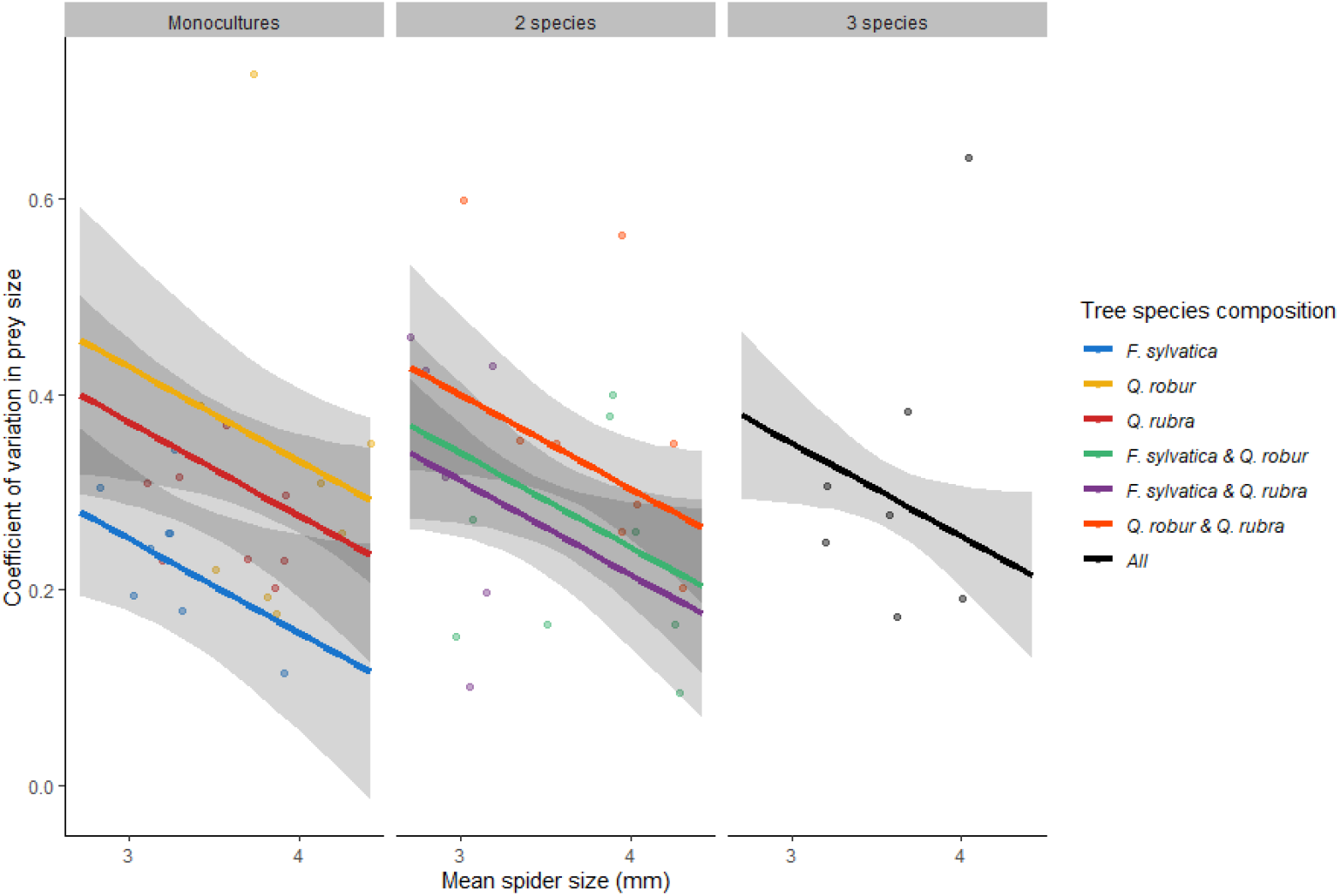
Relationship between the coefficient of variation (CV) for prey size per plot and the mean spider’s cephalothorax size (mm) per plot. Data points are the individual plots (N = 53). Lines with 95% CI are the estimated slopes based on model M_add_. For estimation, edge distance was taken to be the overall average. Colours refer to the tree species composition. In plots with larger spiders, there is a stronger stability in prey size (lower CV). Additive tree species composition effects are present. The strongest stability in prey size is found in monocultures of *F. sylvatica*, and the lowest stability in monoculture of *Q. robur*. In mixtures, the CV is an average of the CV of each individual monoculture contributing to the mixture.

## Discussion

Tree diversity is an important driver of biodiversity in smaller forest fragments in Flanders (Hertzog et al. 2020), but different taxa react in different ways to the specific environmental gradients (Perring et al. 2021). We here use gut-content metabarcoding to show how changes in tree diversity, identity and edge effects affect an important functional biodiversity component, namely trophic interactions between an orb-web spider and its prey. Tree species composition, rather than tree diversity, impacted diet. More specifically, spiders consistently consumed larger prey in monocultures of *F. sylvatica* than in other tree species compositions. Contrary to expectations, we found prey species richness in the spider diet to decrease with spider size, but the proportion of large prey items to increase. Since spiders were largest in monocultures of *Q. robur* and smallest in monocultures of *F. sylvatica*, but also larger closer to the forest edge, size-mediated interactions are prevalent. Plots inhabited by on average larger spiders showed lower individual-variation in prey (size) consumption

Unlike our expectation that plant diversity impacts insect diversity and trophic interactions (Price, 2002; Scherber *et al*., 2010; Rzanny *et al*., 2013), we did not find any effect of tree diversity on the richness or size of prey in the diet. The does not imply that plant diversity cannot have an impact on the diet of other predators. In one of the rare studies that focussed on trophic interactions of a generalist predator in relation to plant diversity, it was shown that the richness of consumed prey in carabid beetles did increase along an experimental grassland diversity gradient (Tiede *et al*., 2016). The general expectation that the diet of a predator contains more prey in prey rich habitat, is based on ecological opportunism (Bison *et al*., 2015). Essentially, a generalist predator’s diet would reflect the diversity of prey available. However, this is contradicted by the idea that the availability of more prey species allows intraspecific specialization on species (Staudacher *et al*., 2018). We found no support for increased specialization in more diverse forest, as the coefficient of variation in prey richness did not depend on tree diversity. It should, however, be made clear that the occurrence of ecological opportunism can neither be confirmed nor dismissed as both processes act at the individual level, and may be levelled out at the population-level. In line with earlier research finding mixtures between *Q. robur* and *F. sylvatica* to harbour the highest (arthropod) diversity (Hertzog et al. 2020), we found this two-species mixture to held a different composition of species in the spiders’ diet, but not richness, compared to the other tree species composition. However, if spider diet did indeed reflect the prey availability, we would expect spider diet in forests with *Q. robur* to have the more species and being composed of larger prey, since *Q. robur* is known to harbour a more diverse arthropod community (Southwood, Wint, Kennedy & Greenwood, 2004). This could not be demonstrated, but we need to emphasise that it remains difficult to confidently conclude whether our findings are pure reflections of the prey availability without its effective quantification, or whether some prey selection is prevalent. Indeed, larger spiders tended to be more selective in their foraging by the consumption of fewer, but larger prey species. Consumption of larger prey is likely needed to achieve their higher energy requirement (Brown *et al*., 2004). We can, however, not attribute the general prey size increases to the selection of the largest most energy efficient prey, as prey larger than one cm did not show a clear relationship with spider size. More-over, our approach does neither allow more quantitative analyses on the relative abundances, consumed biomasses or intraspecific variation in prey size. Prey sizes could only be derived from literature, and while caught prey are adult active flying stages, intraspecific variation may be substantial. Irrespective of spider size, spiders also consumed consistently large prey, and more and larger prey of high gain (> 1cm) in monocultures of *F. sylvatica* than in other tree species composition. Interestingly, monocultures of *F. sylvatica* also hold the smallest spiders.

We found no changes in prey richness or composition in relation to edge proximity. A possible pattern could, however, be masked by the occurrence of larger spiders in proximity of the forest edge. Neither larger spiders close to the edge, nor the absence of larger prey species consumed close to the edge fit the expectation that the warmer forest edges could favour smaller arthropods (Atkinson & Sibly, 1997; Kingsolver & Huey, 2008). Wind is stronger in the forest edges (Schmidt *et al*., 2017). Wind damages webs, which reduces the foraging efficiency and enforces costly web repairs (Tew, Adamson & Hesselberg, 2015). This may select for larger spiders with higher silk production (Vollrath, 1999). It is possible that within this study a stronger effect is overlooked, as edge effects for both biotic and abiotic gradients are generally observed in first few meters from the edge (Murcia, 1995; Schmidt, Jochheim, Kersebaum & Lischeid, 2017; De Smedt *et al*., 2018).

In conclusion, we show that an interception predator like the orb web spider *A. diadematus* is not only generalist in prey capture but also prey consumption. At the individual level, however, the prey spectrum of this generalist predator is linked to body size, with larger spiders foraging on a smaller number of consistently larger prey species. Contrary to expectations from earlier research on insect diversity in forest fragments differing in tree species composition, prey species richness and size are only marginally explained by tree species composition. Edges effects on trophic relations were only indirectly prevalent through differences in spider size.

## Acknowledgements

We thank the research group of Tomas Roslin for providing the workspace for performing the molecular analysis, all the forest owners for their permission to conduct our sampling in their forest, Daan Dekeukeleire for many discussions and Pieter Vantieghem for his field work assistance. This study was financially supported by Concerted Research Actions – Special Research Fund – Ghent University for the UGent GOA project Scaling up Functional Biodiversity Research: from Individuals to Landscapes and Back (TREEWEB) and FWO project G080221N.

## Data accessibility Statement

All data on spider size, plot composition and DNA sequence assemblies are available on DRYAD: https://doi.org/10.5061/dryad.7sqv9s4s3

## Author contributions

IMSL, AM, KV, LL, DB designed the study, EJV assisted IMSL with the molecular analysis, LRH, DB assisted IMSL with the data-analysis. All authors contributed to the writing of the manuscript.

